# The chloroplast ATP synthase redox domain in *Chlamydomonas reinhardtii* attenuates activity regulation as requirement for heterotrophic metabolism in darkness

**DOI:** 10.1101/2022.11.08.515721

**Authors:** Lando Lebok, Felix Buchert

## Abstract

To maintain CO_2_ fixation in the Calvin Benson-Bassham cycle, multi-step regulation of the chloroplast ATP synthase (CF_1_F_o_) is crucial to balance the ATP output of photosynthesis with protection of the apparatus. A well-studied mechanism is thiol modulation; a light/dark regulation through reversible cleavage of a disulfide in the CF_1_F_o_ γ-subunit. The disulfide hampers ATP synthesis and hydrolysis reactions in dark-adapted CF_1_F_o_ from land plants by increasing the required transmembrane electrochemical proton gradient 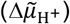. Here, we show in *Chlamydomonas reinhardtii* that algal CF_1_F_o_ is differently regulated in vivo. A specific hairpin structure in the γ-subunit redox domain disconnects activity regulation from disulfide formation in the dark. Electrochromic shift measurements suggested that the hairpin kept wild type CF_1_F_o_ active whereas the enzyme was switched off in algal mutant cells expressing a plant-like hairpin structure. The hairpin segment swap resulted in an elevated 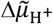 threshold to activate plant-like CF_1_F_o_, increased by ∼1.4 photosystem (PS) I charge separations. The resulting dark-equilibrated 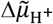 dropped in the mutants by ∼2.7 PSI charge separation equivalents. Photobioreactor experiments showed no phenotypes in autotrophic aerated mutant cultures. In contrast, chlorophyll fluorescence measurements under heterotrophic dark conditions point to a reduced plastoquinone pool in cells with the plant-like CF_1_F_o_ as the result of bioenergetic bottlenecks. Our results suggest that the lifestyle of *Chlamydomonas reinhardtii* requires a specific CF_1_F_o_ dark regulation that partakes in metabolic coupling between the chloroplast and acetate-fueled mitochondria.

**Significance Statement:** The microalga *Chlamydomonas reinhardtii* exhibits a non-classical thiol modulation of the chloroplast ATP synthase for the sake of metabolic flexibility. The redox switch, although established, was functionally disconnected in vivo thanks to a hairpin segment in the γ-subunit redox domain. Dark enzymatic activity was prevented by replacing the algal hairpin segment with the one from land plants, restoring a classical thiol modulation pattern. Thereby, ATP was saved at the expense of thylakoid membrane energization levels in the dark. However, metabolism was impaired upon silencing dark ATPase activity, indicating that a functional disconnect from the redox switch represents an adaptation to different ecological niches.

## Introduction

The chloroplast ATP synthase (CF_1_F_o_) is the major H^+^ gate of the photosynthetic machinery (reviewed in 1, 2). Its driving force is the transmembrane electrochemical proton gradient (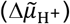) that stems from coupled photosynthetic electron transfer and H^+^ movements across the thylakoid membrane. Therefore, 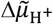 consists of a membrane potential (ΔΨ) and an osmotic component (ΔpH) which both catalyze ATP synthesis (3-6). CF_1_F_o_ consists of two parts (7, 8), a soluble (F_1_with subunits α_3_β_3_γ_1_δ_1_ε_1_) and a transmembrane moiety (F_o_with subunits IV_1_I_1_II_1_III_14_or a_1_b_1_b’c_14_). The electrochemical F_o_motor translocates H^+^ across the membrane through reversible protonations that drive the rotation of the c_14_-ring, which is transmitted to the coupled γ- and ε-subunits. Nucleotide (de)phosphorylation takes place in the chemical F_1_ motor, containing three catalytic binding pockets within the β-subunits that assume different conformations (9). This is due to steric clashes with the asymmetrical, rotating crank structure of the γ-subunit helical termini which push aside a helix-turn-helix element in the β-subunit, the DELSEED motif. Thus, the mechanical coupling of F_1_ and F_o_ reactions facilitate the equilibration of the free energy stored in the phosphorylation potential [ATP]/([ADP][Pi]) and in the 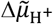, respectively.

With its H^+^-gating function, CF_1_F_o_needs to be fine-tuned for optimal photosynthetic ATP output and NADPH formation efficiency at photosystem I (PSI) to sustain CO_2_ fixation in the Calvin Benson-Bassham cycle. The electrochromic shift (ECS) of photosynthetic pigments has been used extensively to describe CF_1_F_o_ activity states in vivo (10, 11, reviewed in 12). The pigment absorption changes in response to ΔΨ and ECS decay signatures depend on ATP synthesis and ion channel activities. A poor H^+^ conductivity across the membrane would generate an excessive ΔpH that counters photosystem II (PSII) efficiency and stability (13, 14), eventually affecting NADPH and ATP yields. Moreover, excessive pH-dependent nonphotochemical quenching (reviewed in 15) would become restrictive for photochemical productivity. It has also been shown that deregulated CF_1_F_o_ favors large ΔΨ spikes under fluctuating light that damage PSII (16). Therefore, CF_1_F_o_ performance control in concert with other ion channels/antiporters (17, 18) are vital for optimal parsing of 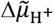 components.

Various environmental factors regulate CF_1_F_o_ activity such as high light, drought, or CO_2_ availability (19-21) but the most well-known CF_1_F_o_ regulation involves “thiol modulation”. A redox-active cysteine couple in the γ-subunit forms a disulfide that is cleaved enzymatically in the light (22-24) and restored in darkness (25). Disulfide cleavage lowers the 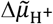 threshold required for the onset of reversible ATP synthase activity (26, 27) and the dithiol formation in low light allows for efficient electron flow via a highly H^+^-conductive CF_1_F_o_. Besides the cysteine couple, a structural determinant for H^+^ conductivity tuning is a hairpin within the γ-redox domain that interacts with the β-subunit DELSEED motif (7, 28). The full scope of thiol modulation is not yet understood as mutants of the redox couple are vital in *Arabidopsis thaliana* (hereafter *A. thaliana*) (29) and *Chlamydomonas reinhardtii* (hereafter *C. reinhardtii*) (30). However, deactivating ATPase activity in the dark via disulfide formation was proposed to serve as an ATP-preserving strategy (31). Yet, in the presence of an inactive CF_1_F_o_, other processes are required to sustain the polarization state of the membrane, i.e., to generate a 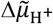 in darkness 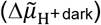. The 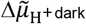 could function on several levels, such as facilitating protein import into the lumen (32) or priming CF_1_F_o_ to surpass its 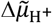 activation threshold at the onset of light. Chlororespiration, which is important for a multitude of processes (reviewed in 33), could fulfil the requirements for sustaining 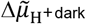. Chlororespiratory electron flow occurs between NAD(P)H and oxygen by tandem action of an NADPH dehydrogenase (NDH) and a plastid terminal oxidase (PTOX), transiently storing the electrons in the plastoquinone pool. NDH activity in *A. thaliana* is electrogenic since it pumps H^+^ into the lumen whereas this is not the case for the Type II NDH in *C. reinhardtii*, NDA2. Therefore, algae were speculated to sustain 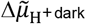 by ATP hydrolysis instead of chlororespiration (33, 34).

Here, ECS assays were used to demonstrate that thiol modulation in *C. reinhardtii* cells is differently functionalized than in land plants, owing to a hairpin structure that allows the alga to energize its thylakoid membranes throughout the night. We also show that algal CF_1_F_o_must remain active in the dark to balance the cellular redox poise via chlororespiration, which strongly depended on the trophic state of the algal cultures.

## Results

### 1. Linking in vivo CF_1_F_o_ activity upon dark adaptation to sequence variants in the γ-subunit redox domain

The thiol switch from *C. reinhardtii* CF_1_F_o_ γ-subunits displays the expected regulatory patterns in vitro and ATP synthesis rates at lower ΔpH were recorded in the reduced dithiol conformation (28). Nevertheless, we noticed under disulfide promoting conditions in dark-adapted photoautotrophic *C. reinhardtii* cultures that the enzyme was different from land plant leaves. A common in vivo method to visualize γ-subunit thiol modulation upon dark adaptation is the use of flash-induced ECS decay kinetics (29, 35). The assumption is to link a fast ECS decay to a highly active enzyme catalyzing ATP synthesis in the presence of a γ-dithiol. Flash-induced ECS measurements were carried out in a collection of land plants and *C. reinhardtii* (Fig. 1A). All samples were dark-adapted for 30 min and, as shown in the inset, the typical three phases of the ECS kinetics were visible (reviewed in 12): The rapid ECS increase directly after the flash was attributed to PSI and PSII charge separation activity (a-phase). This was followed by a ∼10-ms rise in ECS which is related to charge separation in the low-potential chain of the cytochrome *b*_6_*f* complex (b-phase). Eventually, the ECS decay mainly results from ATP synthesis activity through CF_1_F_o_ (c-phase). Clearly, the latter phase was most rapid in dark-adapted *C. reinhardtii* cells. In a next step we tested whether this might be linked to a hairpin loop in the γ-subunit that plays a role in redox regulation of CF_1_F_o_ (7, 28). To this end we followed a segment swapping approach to assess this by genetic engineering of the γ-subunit loop. Fig. 1B shows a partial alignment of the redox domain and underlines the variations within the γ-subunit loop found in species from Fig. 1A. Moreover, the genetic sequence of the γ-subunit construct that we created to mimic a plant-like CF_1_F_o_ in *C. reinhardtii*, termed γloop, is also listed. Fig. 1C provides the spatial reference of the aligned segment and highlights the critical position of the loop within the α_3_β_3_ hexamer (7). It is known that *C. reinhardtii* cells establish a γ-disulfide in the dark (28) and by creating γC233S that lacks one of the redox-active cysteines we observed an accelerated c-phase in the flash ECS experiments (Fig. 1D). Importantly in Fig. 1D, dark-adapted γloop mutants showed a slow ECS decay that resembled land plant kinetics from Fig. 1A. When disulfide formation was prevented in γloop/γC233S the ECS decay was accelerated (Fig. 1D, see SI Appendix, Fig. S1 for biological replicates).

**Figure 1.**
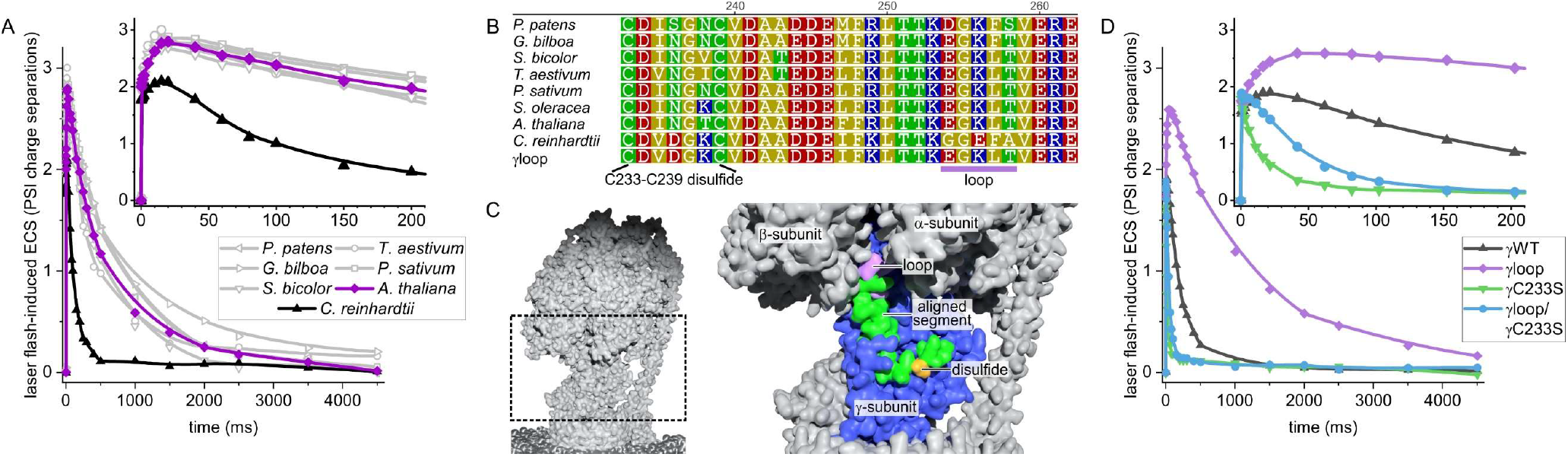
A loop segment in the algal γ-subunit redox domain determines laser flash-induced decay kinetics of the ECS signal upon dark adaptation. (*A*) Decay of ECS kinetics was measured in dark-adapted *C. reinhardtii* and various land plant species. (*B*) The partial γ-subunit redox domain sequence alignment of species from panel *A* highlight the varied loop segment that was exchanged in γloop algal mutants (*C. reinhardtii* numbering on top). (*C*) The rotating loop segment is inserted in between the α_3_β_3_ hexamer, shown here for spinach CF_1_F_o_ (PDB ID: 6FKH). (*D*) CF_1_F_o_ was analyzed in photoautotrophic algal cultures via flash-induced ECS decay measurements. Besides plant-like γloop mutants the analysis also involved two γC233S disulfide mutants in the presence of both loop variations (see SI Appendix, Fig. S1 for biological replicates).

### 2. The natural sequence variant in the algal γ-subunit redox domain determines the dark-equilibrate 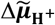

It is known that *C. reinhardtii* establishes the γ-disulfide in darkness which is cleaved upon illumination (28), and we confirmed thiol modulation for γWT and γloop (SI Appendix, Fig. S2 and SI Materials and Methods). The ECS decay in Fig. 1D was driven by single-turnover laser flashes that excite all photosystems, but ECS amplitudes were normalized to PSI. Accordingly, one flash perturbed the dark-equilibrated 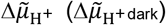 by ∼1.75 PSI charge separations to drive ATP synthesis activity via CF_1_F_o_ (see also SI Appendix, Fig. S1). It is known that a large 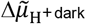 amplitude can override functional CF_1_F_o_ restrictions imposed by the γ-disulfide (26, 27), which might account for the ECS decay differences in Fig. 1D. To address the issue, ECS-based estimations of 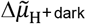 were obtained in those samples using short saturating light pulses to catalyze multiple turnovers in the photosystems and cytochrome *b*_6_*f* complex (Fig. 2A; for details of this paragraph see Material and Methods). Once the membranes were substantially energized by the light pulse, the ECS signals returned to the respective 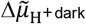 baseline within several seconds in the dark, owing to ATP synthesis activity. The initial rates of ECS generation at the beginning of the light pulse were identical (yellow symbols in Fig. 2A, SI Appendix, Fig. S3), which point to similar photochemical efficiencies after the dark adaptation. However, the ECS amplitude during the pulse varied in the CF_1_F_o_ mutants. The variation stemmed from the fact that the respective baseline 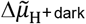 differed. This relationship – the larger the ECS amplitude during the pulse, the lower was 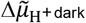 in the sample (27, 36) – is revealed when collapsing 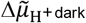 via respiration inhibitors or uncouplers (SI Appendix, Figs. S4A-B). In the absence of inhibitors, the dark-adapted γloop in the representative measurement shown in Fig. 2A produced larger signals during the light pulse (due to lower 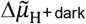), whereas both γC233S and γloop/γC233S produced slightly smaller signals than γWT. We noticed that γC233S did hardly produce a stable ECS plateau during the pulse. The reason for this remains unknown but the ECS drifts were inducible in γWT when lowering the pulse light intensity (SI Appendix, Fig. S4C). However, the ECS plateau level at the end of the pulse was not used for the following ECS-based 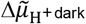 estimations.

**Figure 2.**
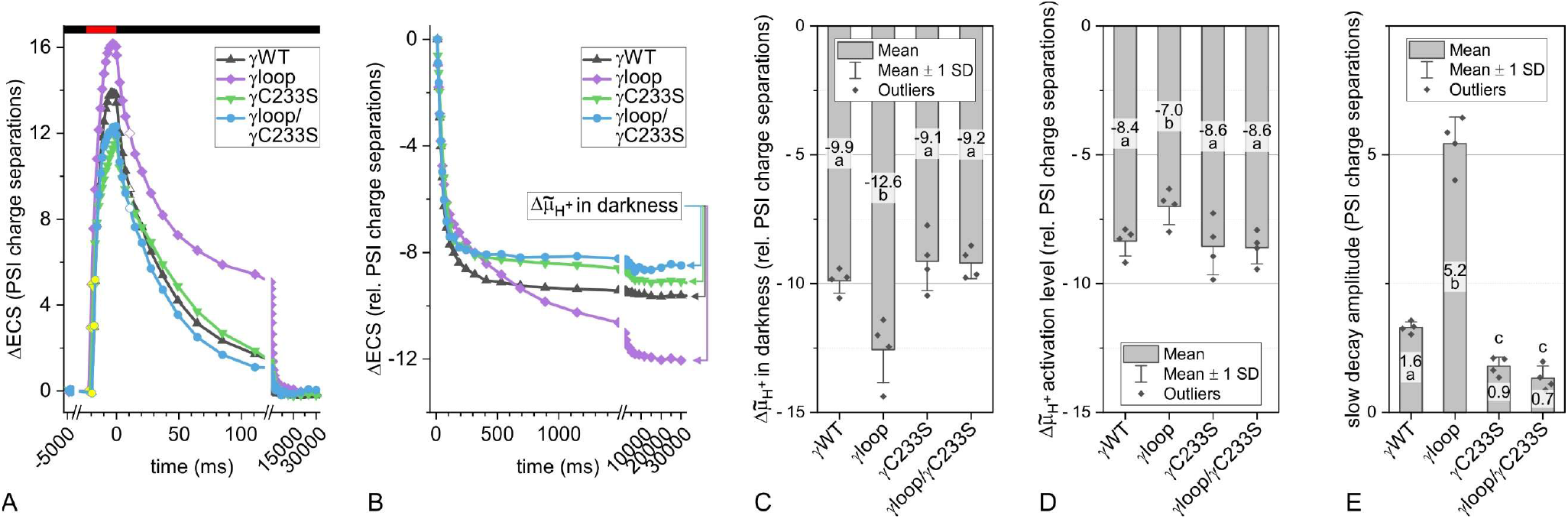
Optical measurements of the 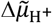 in the dark reveal lower values in the CF_1_F_o_ γloop mutant due to its elevated activation threshold. (*A*) The increase of the ECS by a light pulse (red bar) as well as the ECS decay in dark (black bar) is shown in samples from Fig. 1D. Linear initial ECS generation was quantified from yellow symbols (see SI Appendix, Fig. S3). To deconvolute the variable ECS baseline level, the technical ECS reference at 10 ms after the pulse (white symbols) was used. (*B*) This reference was set to 0 for expressing the individual 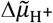 in darkness and comparing the multiphasic ECS decay. (*C*) The averaged 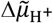 in darkness is shown (*N* = 4 ± SD, One-Way ANOVA/Fisher-LSD, *P*<0.05). (*D*) ECS decay kinetics from panel *B* were fit with a two-exponential (see SI Appendix, Fig. S5 for biological replicates) and the averaged 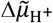 activation level is shown that separated the fast from the slow decay phase. (*E*) The averaged amplitudes of the slow ECS decay phase are shown.

Instead, the obscure 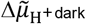 baseline was estimated by referencing it to ECS_10ms_, i.e., the measured ECS at 10 ms after the pulse that was arbitrarily defined as 0 (27, 36). Accordingly, the 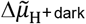 was expressed as negative PSI charge separation equivalents (Fig. 2B). The kinetics comparison further revealed that the ECS decay after the light pulse was biphasic in γloop and virtually monophasic in the other strains. When treating the ECS data to quantify 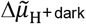 statistically, -9.9 PSI charge separation equivalents were measured in γWT, whereas the levels were insignificantly raised in γC233S and γloop/γC233S (Fig. 2C). The 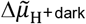 of -12.6 PSI charge separation equivalents was significantly decreased in γloop, and the difference disappeared upon uncoupler treatment where γWT and γloop showed very low 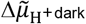 values (SI Appendix, Fig. S4D).

The photosynthetic membranes in Fig. 2B transitioned from ECS_10ms_ (strongly disequilibrated) to 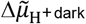 (equilibrated) in a fashion that can be described with a two-exponential function (see SI Appendix, Fig. S5 for fitted biological replicates). Accordingly, ECS decay amplitudes were determined of the fast phase, produced shortly after the pulse, and the slow phase, produced when approaching the respective 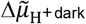. Both ECS decay phases represent fast and slow activity states of the CF_1_F_o_ (37, 38). The fast phase amplitude represents the 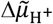 activation threshold level that, once put in relation to ECS_10ms_, is shown in Fig. 2D. Above the 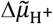 activation level of -8.4 PSI charge separation equivalents the fast CF_1_F_o_ was established in γWT. Very similar thresholds were determined for γC233S and γloop/γC233S. However, the active γloop CF_1_F_o_ was established at a significantly higher level of -7 PSI charge separation equivalents.

### 3. The dark activity of algal CF_1_F_o_ is entangled with metabolic flux control

Despite the differences in darkness (Fig. 2), the less active γloop CF_1_F_o_ was not restricting photosynthetic electron transfer nor PSI reduction during the induction phase of light-adapted cells, suggesting that a limiting ΔpH was not generated under those conditions in the light (SI Appendix, Fig. S6 and SI Materials and Methods). To get physiological insights on why the algal redox domain might have been modified, we investigated in photobioreactor experiments how *C. reinhardtii* performs under diurnal cycles and extended dark periods (Fig. 3A). The bioreactors were incubated with synchronized cell cultures and the 16-h photoperiod was carried over for two cycles. We did not detect substantial differences under photoautotrophic conditions, including the subsequent dark period (black and violet in Fig. 3A; see SI Appendix, Fig. S7 for replicates with independent transformant lines). The OD_700nm_ was set to 0.1 during the first night and the increase rate varied to some extent between transformants but did not substantially differ over the time course of the experiment. Differences were seen for the PSII quantum yields but the trend was not confirmed in the other transformants. The PSII quantum yields were inversely proportional to the light intensity and the lowest fractions of open reaction centers ranged from 0.15-0.35 at the peaking 250 μmol photons m^-2^ s^-1^. However, during the dark phases the PSII quantum yields (here, F_v_/F_m_) remained stable at ∼0.75 (Fig. 3A). We also examined the γWT and γloop under photoheterotrophic light/dark regimes in the presence of acetate (blue and orange in Fig. 3A, SI Appendix, Fig. S7). Here, the fraction of open PSII in the peaking light intensities was slightly increased compared to acetate-free conditions, ranging from 0.2-0.45, and the steep decline of PSII quantum yields occurred at higher light intensities. Increased PSII quantum yields in the presence of acetate are linked to diminished energy-dependent de-excitation processes. The latter could stem from diminished expression of light harvesting complex (LHC) stress-related3 protein, LHCSR3 (39), which was not further explored here. Besides LHCSR3-associated processes, the transformants in the photobioreactor showed a slight variation of their chlorophyll content (SI Appendix, Fig. S8). Smaller PSII antenna sizes as a result can therefore not be ruled out to favor higher PSII quantum yields (40) in some transformant lines. Moreover, the presence of acetate caused a mild but significant decrease in F_v_/F_m_ under extended dark conditions in γWT – although this was not the case in all transformants (cf. black and blue in Fig. 3A and SI Appendix, Fig. S7). Importantly, a consistent effect of acetate that also differed from γWT was seen in all γloop transformants (cf. violet and orange in Fig. 3A and SI Appendix, Fig. S7): The F_v_/F_m_ declined significantly in the mutant in an oscillating pattern which loosely aligned with the previous photoperiods. However, this effect phased out during the second day in the dark and sporadic increases of F_v_/F_m_ were observed. F_v_/F_m_ recovery in γloop coincided with cell growth arrest, which we attribute to depletion of acetate in the medium. The time of growth cessation varied in the transformants, but the F_v_/F_m_ reversion was exclusively seen in γloop. We also noticed that the onset of declining F_v_/F_m_ and the reversion of the effect coincided with the respective rise and relaxation of F_0_, the basal chlorophyll fluorescence (SI Appendix, Fig. S9). This points to a reversible reduction of the plastoquinone pool (41) and the metabolic F_v_/F_m_ signature in heterotrophic γloop prompted us to investigate chlororespiration mutants in this experimental context. Indeed, a virtually coinciding recovery of low F_v_/F_m_ and a stagnation of cell growth upon acetate depletion was also observed in *ptox2* mutants (Fig. 3B, SI Appendix, Fig. S10). The reduction of the plastoquinone pool in the dark in the absence of the major plastoquinol oxidase PTOX2 strongly depended on the availability of fixed carbon in the form of acetate (cf. green and olive in Fig. 3B and SI Appendix, Fig. S11). The reduced plastoquinone pool triggers state transitions upon STT7 kinase activation and migration of phosphorylated LHCII towards PSI (reviewed in 42), lowering F_v_/F_m_. The extent of F_v_/F_m_ recovery upon acetate depletion, however, was less pronounced in γloop vs. *ptox2*.

**Figure 3.**
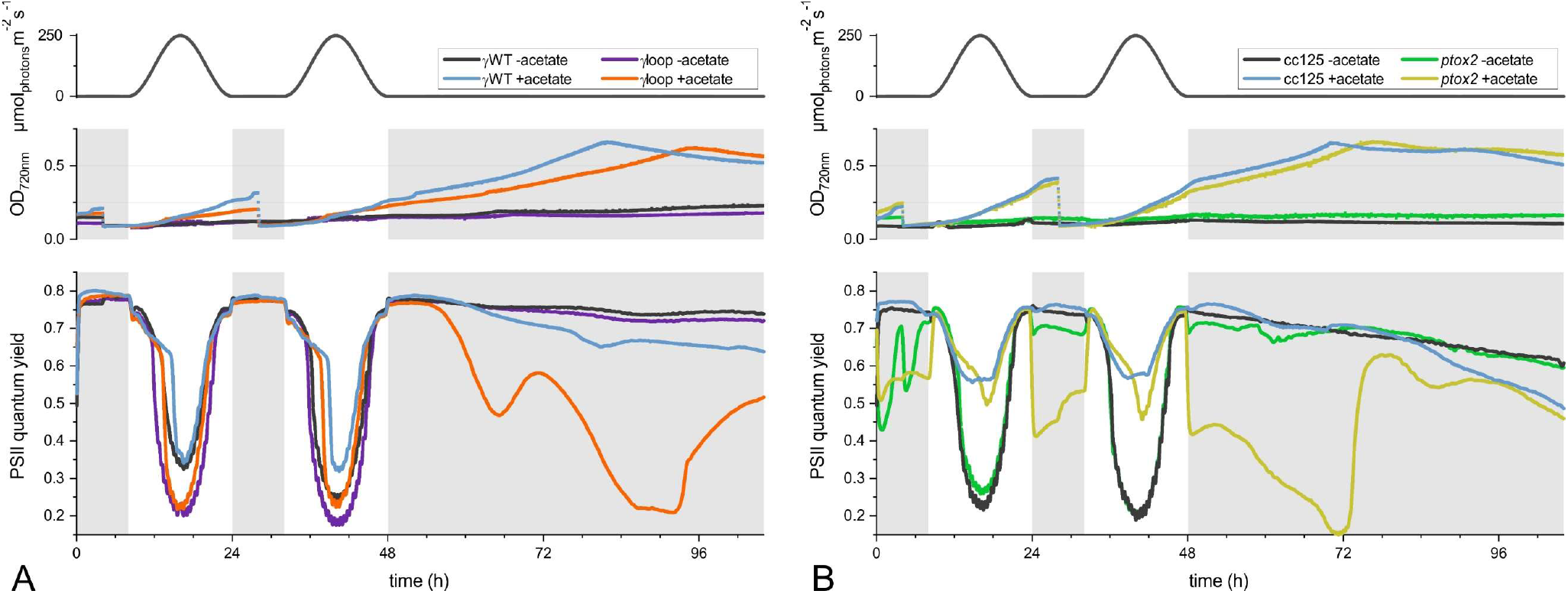
Algal photobioreactor experiments show two light cycles and exposure to extended darkness. (*A*) The OD_720nm_ turbidity, adjusted during the first night cycle (and second in the presence of acetate), as well as the PSII quantum yields are shown for CF_1_F_o_ variants γWT and γloop mutant at a given light intensity. (*B*) The same experiment is shown for CC-125 and *ptox2* (see SI Appendix, Figs. S7 and S10 for replicates).

## Discussion

### Escaping thiol modulated activity regulation due to the natural sequence variant in the algal γ-subunit redox domain

The entanglement of thiol modulation – i.e., the reversible γ-disulfide cleavage – and the 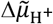 required to drive ATP synthesis has been consistently shown in liposome reconstitution assays from spinach (26) and *C. reinhardtii* CF_1_F_o_ (28), as well as in *A. thaliana* leaves (27). It is consensus that the oxidized γ-disulfide conformation requires a higher 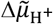 to be active. However, published data suggests that thiol modulation by itself is less efficiently realized in *C. reinhardtii*. Hisabori and co-workers showed in in vitro experiments from *A. thaliana* that the half-time of γ-disulfide cleavage by recombinant *f*-type thioredoxins is about 1 min (43). The group has also conducted these experiments in the *C. reinhardtii* context, showing that half reduction took about 7 min (28). These studies (28, 43) also report a much slower γ-disulfide cleavage in dark-adapted algal cells vs. *A. thaliana* leaves, using the same light intensity (half reduction in vivo ∼65 s vs. ∼10 s in *A. thaliana*). However, unlike the high AMS labeling efficiency for the *A. thaliana* γ-subunit in the light as a full γ-disulfide cleavage indicator, the labeling efficiency was somewhat lower in the previous algal study (28), which resembled our findings (SI Appendix, Fig. S2). This raises the question why the algal system tolerates a slower, less efficient redox tuning of the chloroplast ATP synthase?

Here, we have demonstrated in vivo that the typical thiol modulation concept has been attenuated in dark-adapted *C. reinhardtii* cells via its intrinsic hairpin segment from Figs. 1B and 1C. The conclusion is based on the observation that the ECS kinetics of dark-adapted γloop algae resemble those from spinach and *A. thaliana* (27, 35, 36). In contrast, the 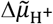 activation level of algal CF_1_F_o_ remained relatively low, whereas it was raised by 1.4 PSI charge separations in the plant-like γloop (Fig. 2D). The mutant 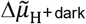 dropped by 2.7 PSI charge separation equivalents (Fig. 2C) as a result of a larger pool of slowly active CF_1_F_o_. The latter produced a larger amplitude of the slowly decaying ECS in Fig. 2E which also explains the flash kinetics in Fig. 1D. Therein, both reaction centers produced less than 2 PSI charge separations after the flash as the driving force for ATP synthesis. Only the slow γloop CF_1_F_o_ operated far below its 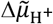 activation threshold, unlike γWT and both γC233S variants. One possible explanation of the lower 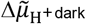 in γloop could be that ATPase-driven lumen acidification in the dark is not sufficient to counter passive ion leaks. Trans-thylakoid H^+^ pumping that relied on the free energy in the phosphorylation potential [ATP]/([ADP][Pi]) was also depending on mitochondrial activity in the dark (SI Appendix, Fig. S4A). Yet, γloop was inactive despite the available ATP which might relate to a transient inhibition by tightly bound Mg-ADP that is favored under disulfide conditions (44, 45). The affinity of *C. reinhardtii* wild type CF_1_F_o_ for ADP is substantially lower than in spinach (46) and it remains to be tested whether this is due to the hairpin segment. Likewise, future efforts of increasing the available stromal ATP might yield an active γloop in the dark, e.g., through ATP import engineering (see next Section). From a mechanistic view, it is possible that the wild type algal loop segment attenuated the rotational constraints despite the γ-disulfide (Fig. 1B, underlined). The involved salt bridges between the critical arginine of the stationary βDELSEED loop and the γ-glutamate position 256 and were less stable (but not in γloop), thus attenuating the proposed chock that blocks rotation (7).

### Entangling metabolic fluxes with dark activity of C. reinhardtii CF_1_F_o_

Based on the photobioreactor analyses, the plastoquinone pool appeared to be a dead-end in γloop (SI Appendix, Fig. S9). Fig. 4 summarizes our current hypotheses of misregulated metabolic pathways in the mutant, all of which remain subject of active research. Although other processes may be relevant, such as starch metabolism or protein import, the current dataset suggests that chlororespiration is affected in γloop after silencing dark ATPase activity. Chlororespiration (reviewed in 33) may assist in membrane polarization in vascular plants where NDH is electrogenic and dark ATPase activity is low. Contrary, dark ATPase activity via CF_1_F_o_ may be a chlororespiratory prerequisite when a non-electrogenic NDH is involved, such as *C. reinhardtii* NDA2 (47). When there is an excess of ATP in the cytosol in the dark, heterotrophic *C. reinhardtii* cells import ATP into the chloroplast at the cost of co-importing NAD(P)H (reviewed in 48). In support of previous work on algal chlororespiration (49, 50), acetate-fueled mitochondrial respiration in Fig. 3 delivered the stromal electrons that were transiently stored in the plastoquinone pool in dark aerated cultures. Accordingly, chlororespiratory oxidation of the plastoquinone pool might sustain cytosolic ATP import. Various cellular signals might be disturbed upon silencing ATPase activity in γloop (Fig. 4), producing chlorophyll fluorescence patterns similar to *ptox2*. Unlike in γloop, the F_v_/F_m_ in *ptox2* nearly returned to the reference strain level upon acetate consumption. This might be linked to PTOX1-dependent plastoquinone pool oxidation (SI Appendix, Fig. S11). If chlororespiration was indeed impaired in γloop, it might affect both PTOX1 and PTOX2. Unlike in dark-adapted *C. reinhardtii* wild type where PTOX2 cofractionates with membranes (50), the tethering of PTOX from vascular plants is facilitated upon illumination (51, 52). The authors showed that light-induced 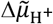 plays a role in PTOX binding to the plant thylakoid membrane. It remains to be tested whether the specific 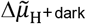 in γloop influences the amounts of soluble PTOX, which lack access to the plastoquinone pool. Thiol modulation of a C-terminal disulfide in PTOX that facilitates reversible dimerization in *A. thaliana* is missing in both PTOX isoforms from *C. reinhardtii* (53). Instead, when aligned with mitochondrial alternative oxidase (AOX), both algal PTOX isoforms harbor a cysteine in vicinity to the dubbed Cys-II, the latter playing a role in redox and metabolic AOX activity regulation (54). Whether factors like ATP, NADPH or other stromal metabolites in γloop have a regulatory effect on PTOX function remains to be tested.

**Figure 4.**
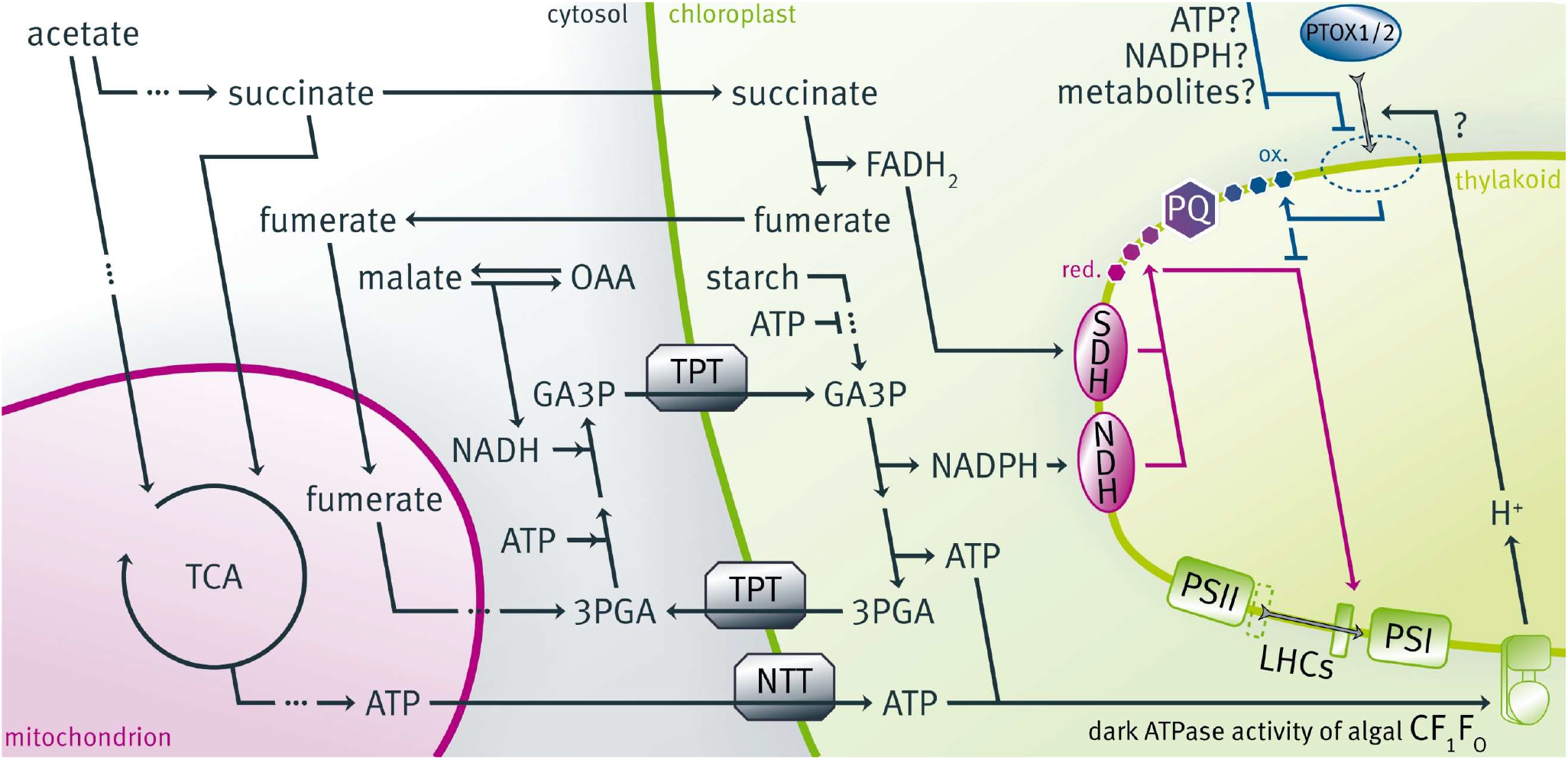
A tentative model is shown of acetate-dependent plastoquinone pool reduction in the absence of dark ATPase activity. Stromal and/or trans-thylakoid determinants might affect PTOX1 and PTOX2 activities in the γloop variant of CF_1_F_o_. The selected metabolic fluxes in *C. reinhardtii* are reviewed in Ref. 48. 3PGA: 3-phosphoglycerate; GA3P: glyceraldehyde 3-phosphate; LHC, light harvesting complex; NDH, NADPH dehydrogenase; NTT, nucleotide triphosphate transporter; OAA, oxaloacetate; PQ, plastoquinone/plastoquinol; PSI/PSII, photosystem I/II; PTOX, plastid terminal oxidase; SDH, succinate dehydrogenase; TPT, triose phosphate/phosphate translocator.

## Materials and Methods

### Transformation and growth of *C. reinhardtii* strains

The transformation plasmid was published previously (19) and is based on the pPEARL expression vector (GenBank: KU531882.1). The *ATPC* CDS was cloned as a *Bam*HI/*Eco*RI fragment using ATPC.F forward (5’GTTCGAATTCATGGCCGCTATGCTCGCC) and reverse ATPC.R primers (5’GCCAGGATCCTTAGCCCGAGGTGGCGGCG). The hairpin that encodes the amino acids 254GGEFA^258^ (*C. reinhardtii* numbering without acknowledging the *ATPC* transit peptide) was mutated using abutting primers hp.F (5’C<u>A</u>AG<u>C</u>T<u>GA</u>CCGTGGAGCGCGAGAAGACC) and hp.R (5’CC<u>CT</u>CCTTGGTGGTCAGCTTGAAGATCTC), indicating underscored mismatches. The 323-bp PCR fragments of ATPC.R/hp2.F and the 775-bp fragment of ATPC.F/hp2.R were ligated, followed by another round of PCR using primers ATPC.F/ATPC.R. The redox-active cysteine couple was disrupted by mutating one of the cysteines at position 203 to serine, using the CS.F (5’CCCATGGGCGAGCTGT<u>CG</u>GACGTGGACGGCAAG). The 406-bp PCR fragment of CS.F/ATPC.R was used in a megaprimer approach (55) to yield the mutated *ATPC* gene in combination with oligos ATPC.F/ATPC.R.

The recipient strain for nuclear transformation did not accumulate *ATPC* transcripts (56) and was cultured on acetate-supplemented TAP medium (57) due to lack of phototrophic growth. Transformants were obtained by electroporation (58) using 25 μF and 1000 V cm^−1^. All successful transformants displayed phototrophic growth and were cultured under diurnal 16-h photoperiods of 50 μmol photons m^-2^ s^-1^ at 23°C on agar-supplemented and liquid Tris-minimal medium (TP) in the absence of acetate (57). Liquid cultures in the presence and absence of acetate were bubbled with sterile air and kept in the exponential phase (0.5–3 × 10^6^ cells ml^-1^) at 23°C by regular dilutions. Air bubbling ensured CO_2_ supply for the Calvin-Benson-Bassham cycle. The experiments in Fig. 1 show the CC-5101 wild type strain. The CC-125 reference strain was used for experiments with *ptox2* mutants (50, 59).

Synchronized cultures (grown under 16-h photoperiods at 60 μmol photons m^-2^ s^-1^) were used when incubating 400 × 10^6^ cells in 400-mL culture volume of TP/TAP in a FMT 150 photobioreactor vessel supplied with sterile air (Photon Systems Instruments, Czech Republic). The culture was diluted to OD_700_ = 0.1 before two consecutive 16-h photoperiods (sigmoidal light 0-250 μmol photons m^-2^ s^-1^ white LEDs) that were followed by more than 48-h darkness at 23°C. TAP cultures were diluted again to OD_700_ = 0.1 after the first photoperiod. Chlorophyll fluorescence measurements were excited with blue LEDs to determine, via readouts of basal (F_0_, F_s_) and maximal fluorescence (F_m_, F_m_’), PSII quantum yields in the dark (F_m_-F_0_)/F_m_, also referred to as F_v_/F_m_, as well as PSII quantum yields in the light (F_m_’-F_s_)/F_m_’.

### Cultivation of land plants

The *Arabidopsis thaliana* (Columbia ecotype) and *Pisum sativum* var. *arvense* were cultivated on soil at 23°C under diurnal 16-h photoperiods of 150 μmol photons m^-2^ s^-1^. *Physcomitrium patens* Gransden wild type was grown at 23°C on PpNO_3_ minimum medium (60) under diurnal 16-h photoperiods of 80 μmol photons m^-2^ s^-1^. Leaves from *Ginkgo biloba, Sorghum bicolor*, and *Triticum aestivum* were obtained from the Botanical Garden of the University of Münster on August 1, 2018 (cloudy, 24°C).

### In vivo spectroscopy

The electrochromic shift (ECS) of photosynthetic pigments was used to study the ΔΨ, the electric 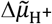 component (10, 11), by measuring the ΔI/I difference at wavelengths 520 nm – 546 nm in a Joliot-type spectrophotometer (JTS-10, Biologic, France, and JTS-150, Spectrologix, USA). The JTS-10 measurements in Fig. 1A used white pulsed LED detection light that was passed through respective interference filters (FWHM: 20 nm). In the other ECS measurements, the JTS-150 pulsed detection light originated from LEDs peaking at 520 and 546 nm, respectively. The light-detecting diodes were protected from scattered actinic light by 3-mm BG39 filters (Schott, Mainz, Germany). Illumination of the samples was interrupted by short dark intervals (250 μs) during which 10-μs detecting pulses were placed after 200 μs. All shown kinetics were obtained in the presence of photosystem (PS) II activity but for comparison reasons the ECS signals were normalized to PSI, i.e., the ΔI/I (520 nm – 546 nm) produced by 1 PSI charge separation in a separate calibration measurement (reviewed in 12). To do so, a saturating 6-ns laser flash was delivered at 700 nm (Q-switched Nd:YAG, Continuum, USA) in the presence of 1 mM hydroxylamine and 10 μM 3-(3,4-dichlorophenyl)-1,1-dimethylurea to inhibit PSII in the algal samples (see SI Appendix, Fig. S12 for algal PSI-related ΔI/I comparison per chlorophyll). In Fig. 1A, it was assumed that land plant PSI and PSII equally contributed to the fast ECS generation upon a laser flash, therefore the flash-induced ΔI/I before the first detection at ∼500-μs was equal to 2 PSI charge separations.

Short saturating light pulses with a duration of several ms were used to drive multiple turnovers of the entire photosynthetic chain until a plateau-like ΔΨ was reached (27, 36). Extending the pulse beyond that point would result in ECS inversion due to ΔpH formation which was avoided by switching off the light. The following ΔΨ consumption was CF_1_F_o_-related and detected as ECS signals returning to the baseline. The latter corresponds to the dark-equilibrated 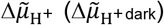, which was not affected by the light perturbations when spacing repetitive measurements by at least 2 min darkness. The ECS signal during a pulse vary in amplitude depending on (*i*) technical and (*ii*) physiological parameters (27, 36). (*i*) The ECS peak amplitude increases with pulse light intensity, which was provided by 630-nm LEDs set to 600 mA in the main text. The same relationship holds for the ECS decay amplitude during the first ∼10 ms after the pulse. However, the subsequent signal amplitude decaying from 10 ms darkness to the baseline 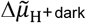 were independent from the light intensity (SI Appendix, Fig. S4C; for the dependency of this amplitude though, see (*ii*) below). Thus, the measured ECS at 10 ms after the pulse (ECS_10ms_) marks a technical reference for light-induced ΔΨ changes with respect to 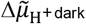. (*ii*) The absolute value of the 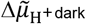 is obscure and a result of cellular physiology. This can be demonstrated by collapsing 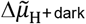 upon treatments with respiration inhibitors or uncouplers (SI Appendix, Figs. S4A-B, S4D) which produced larger pulse-induced signals between 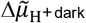 (baseline) and ECS_10ms_ (technical reference). To circumvent this physiological obstruction, the unknown 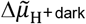 was expressed as negative units relative to the technical reference by setting ECS_10ms_ to 0 PSI charge separations. We resort to this representation in the main text since the alternative approach (measure non-inhibited samples upon dark incubation first, then determine absolute 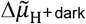 upon inhibitor/uncoupler treatment) is time-consuming and prone to variable metabolic coupling between energy transducing organelles (27, 61).

The decay kinetics between ECS_10ms_ and 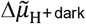 was quantified using the two-exponential function 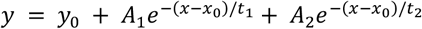. Using OriginPro software, *x*_0_ was fixed to 10 ms, the calculated *y*_0_closely resembled 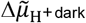, and *A*_l_ (*t*_l_) and *A*_2_ (*t*_2_) correspond to the amplitudes (decay constant) of fast and slow decay phase, respectively (for replicate kinetics and fitted curves see SI Appendix, Fig. S5). The function describes two different CF_l_F_o_ activity states (37, 38): The first one was activated by 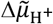 provided by the pulse, whereas the second one was not. Fig. 2D shows −*A*_l_ which marked the 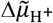 threshold with respect to ECS_10ms_ under which CF_l_F_o_ transitioned to a less active form.

## Acknowledgments

We thank Prof. Michael Hippler for helpful discussions. F.B. acknowledges Deutsche Forschungsgemeinschaft (DFG, German Research Foundation) – 461765884, 507704013.

## Notes

### Competing Interest Statement

The authors have declared no competing interest.

### Summary of Updates

This is a revision of the initial manuscript. The major conclusions were not altered; the data deconvolution was slightly modified. We extended the analysis using biochemical thiol labeling and photobioreactor experiments for photosynthesis measurements, including a chlororespiration mutant.

