## Supplemental Files (Materials and Methods, Figures S1-S12) for "The chloroplast ATP synthase redox domain in *Chlamydomonas reinhardtii* attenuates activity regulation as requirement for heterotrophic metabolism in darkness"

### Supporting Materials and Methods

**In vivo spectroscopy.** The data in Fig. S6 was measured in a JTS-150 (Spectrologix, USA) using differential ΔI/I signals. The ECS ($\Delta I/I_{520nm\text{-}546nm}$; 3-mm BG39 filters, Schott, Germany) was normalized to 1 PSI charge separation (reviewed in 1). To do so, a saturating 6-ns laser flash was delivered at 700 nm (Q-switched Nd:YAG, Continuum, USA) in the presence of 2 mM hydroxylamine and 20 µM 3-(3,4-dichlorophenyl)-1,1-dimethylurea to inhibit PSII in the algal samples. Figs. S6A-B show photochemical rates that were calculated through the change of slope of the ECS signal at the offset of light, based on the Dark Interval Relaxation Kinetics approach (2, 3). Four consecutive detections in the light, each spaced by ~1 ms, were followed by the same series of detection in a ~5ms dark window (4, 5). Illumination of the samples was interrupted by short dark intervals (250 µs) during which 10-µs detecting pulses were placed after 200 µs. The algal samples were adapted to cycles of 2.5 s light (550 µmol photons $m^{-2}$ $s^{-1}$ via 630-nm LEDs) and 2.5 s dark, and 6 cycles were averaged for calculations. Using the same adapted samples, Fig. S6C shows redox populations of $P_{700}$, the primary electron donor of PSI (6). The optical signals ($\Delta I/I_{705nm\text{-}740nm}$; 3-mm RG695 filters, Schott, Germany) determined the fractions of photooxidizable (by a 25 ms saturating pulse), pre-oxidized in actinic light and non-photooxidizable $P_{700}$.

#### Thiol labeling

Labeling of free cysteines with 4-acetamido-4'-maleimidylstilbene-2,2'-disulfonic acid (AMS, Invitrogen) was slightly modified (7). Briefly, photoautotrophically grown cells were harvested in the mid-log phase and resuspended in Tris-minimal medium at $5 \times 10^6$ cells per ml. Dark adaptation (30 min, shaking at 100 rpm) was followed by 10-min illumination at 150 and 500 µmol photons $m^{-2}$ $s^{-1}$, respectively. Cells were precipitated in methanol/chloroform (8) and AMS labeling (2 mM) was carried out in darkness for 1 h at 37°C, loading proteins equivalent to $5 \times 10^6$ cells per lane for a non-reducing SDS-PAGE . Immunostaining via anti-ATPC and anti-AtpB (Agrisera, Sweden) was optionally quantified with Image Lab Software (Bio-Rad, USA).

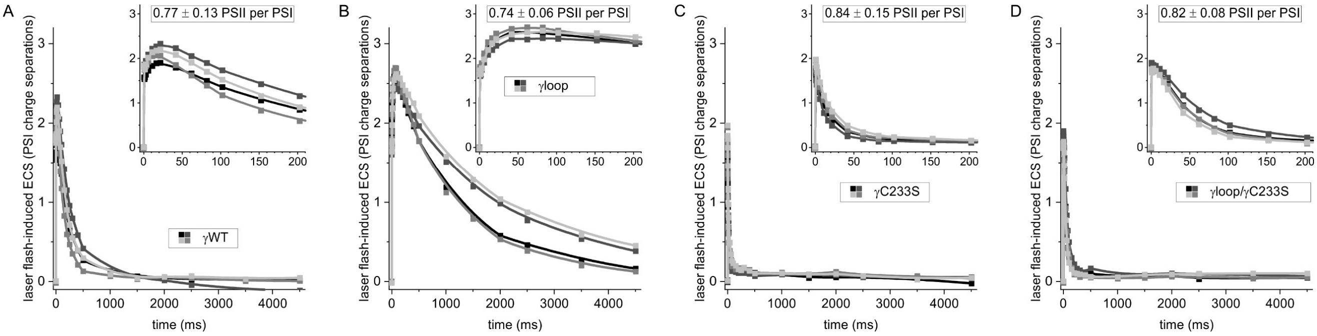


Fig. S1. A loop segment in the algal γ-subunit redox domain determines laser flash-induced decay kinetics of the ECS signal upon dark adaptation. Four biological replicates of (*A*) γWT, (*B*) γloop, (*C*) γC233S and (*D*) γloop/γC233S are shown. The panel insets also show averaged PSI:PSII ratios obtained from the rapid ECS rise after the flash at *t* = 0 ms, which produced an amplitude of 1 PSI charge separation when PSII was inhibited during a separate measurement of the same samples (kinetics not shown). See Main Manuscript Figure 1 as well as Materials and Methods for more information.

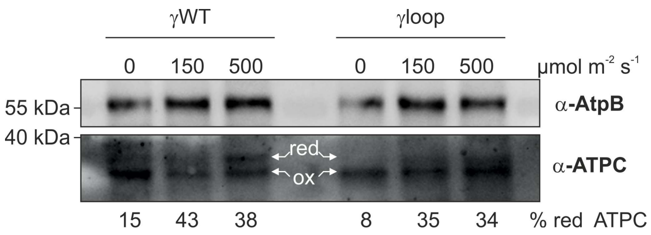


**Fig. S2.** The light-dependent γ-disulfide cleavage is shown. The redox active cysteines of the reduced (red) but not the oxidized (ox) ATPC polypeptide ($CF_1F_o$ γ-subunit) were labeled by 4-acetamido-4'-maleimidylstilbene-2,2'-disulfonic acid, resulting in a migration shift during non-reducing SDS-PAGE. The AtpB signal ($CF_1F_o$ β-subunit) was used as loading control.

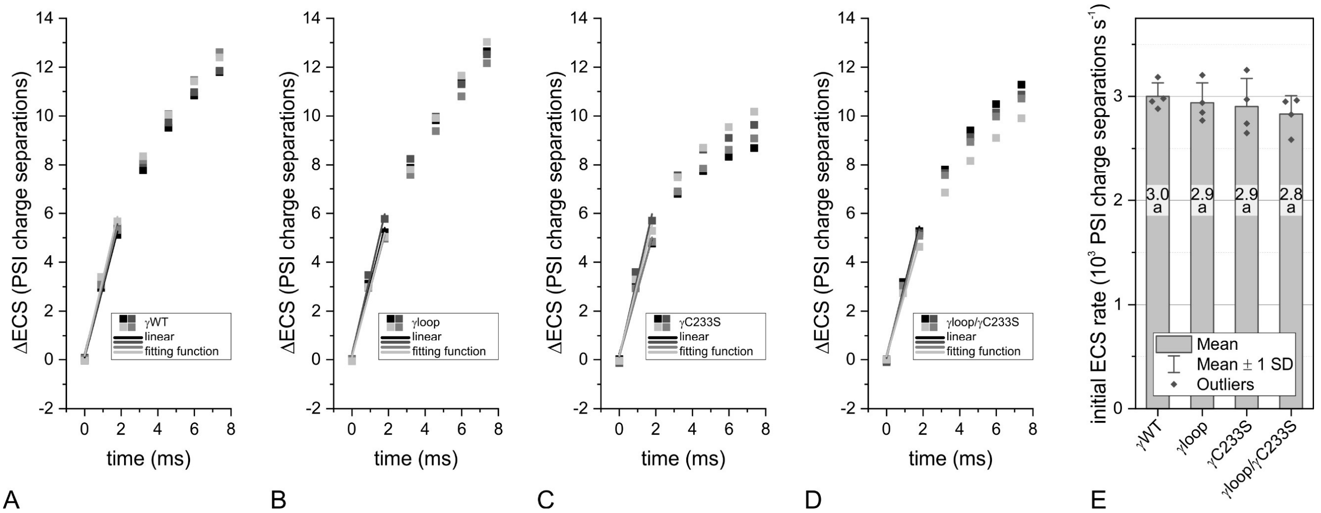


**Fig. S3.** The initial charge separation rates during the saturating pulse are shown. Four biological replicates of (*A*) γWT, (*B*) γloop, (*C*) γC233S and (*D*) γloop/γC233S have been analyzed. (*E*) The linear rise during the first ~2 ms of light was quantified ($N$ = 4 ± SD, One-Way ANOVA/Fisher-LSD, $P$ > 0.05).

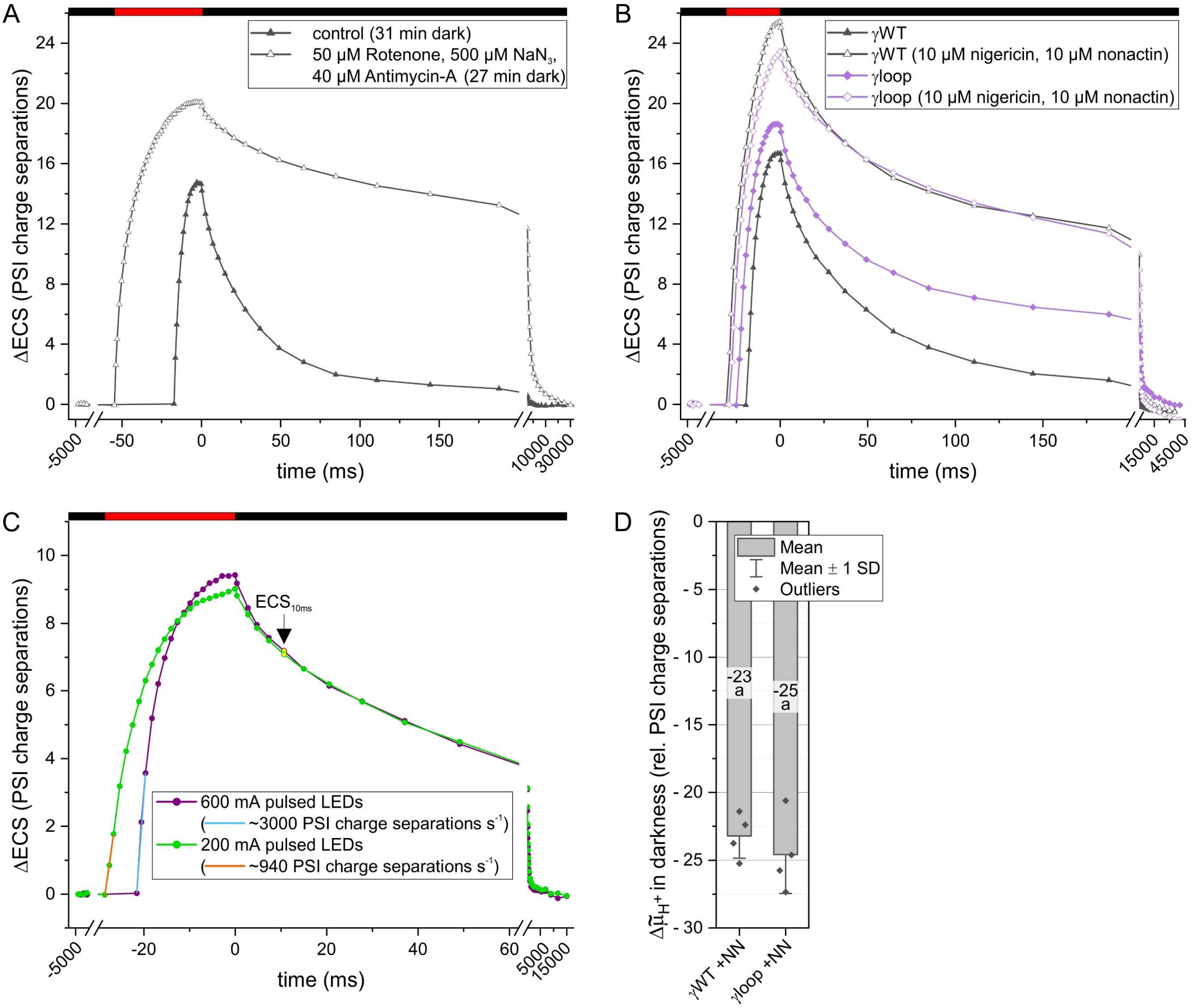


**Fig. S4.** ECS pulse measurements are an optical assay to estimate the $\Delta\tilde{\mu}_{H^+}$ in the dark. (*A*) A cocktail of mitochondrial respiration inhibitors (9) and (*B*) membrane uncoupler treatments result in larger ECS signals generated during the saturating light pulse (red bars). (*C*) ECS signal amplitudes depend on the pulsed light intensity. The ECS value at 10 ms darkness ($ECS_{10ms}$, arrow) was used as technical reference. (*D*) The $\Delta\tilde{\mu}_{H^+}$ in the dark collapsed in the presence of the $H^+/K^+$ exchanger nigericin and the ionophore nonactin (+NN), using conditions as in panel *B* ($N = 4 \pm$ SD, One-Way ANOVA/Fisher-LSD, $P > 0.05$). See Main Manuscript Figure 2C for control conditions.

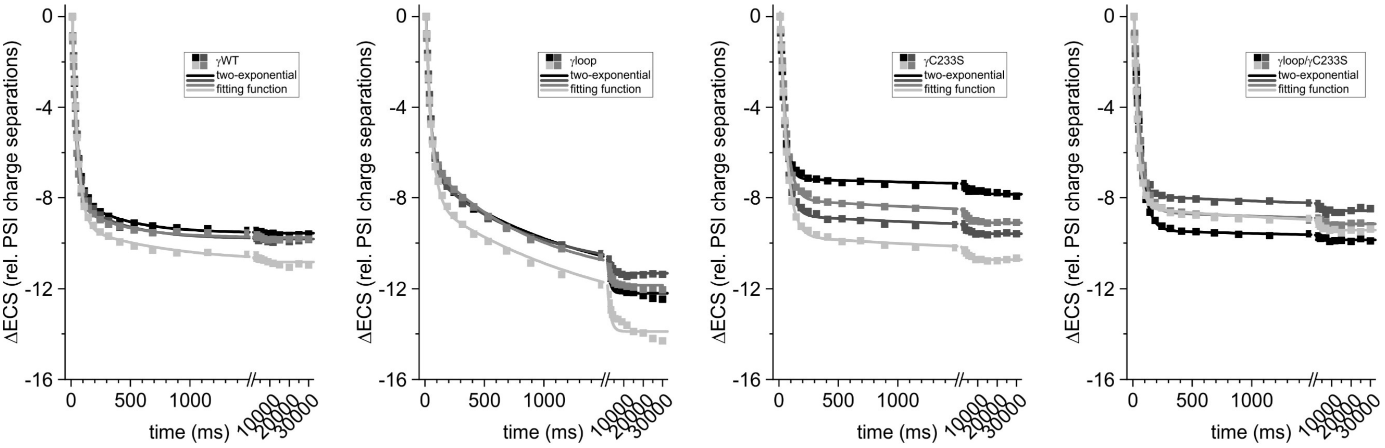


A B C D

**Fig. S5.** Two-exponential fits of pulse-induced ECS decay measurements are shown. The raw kinetics of biological replicates from (*A*) γWT, (*B*) γloop, (*C*) γC233S, and (*D*) γloop/γC233S were referenced to $ECS_{10ms}$. See Fig. S4C and Material and Methods in Main Manuscript for details.

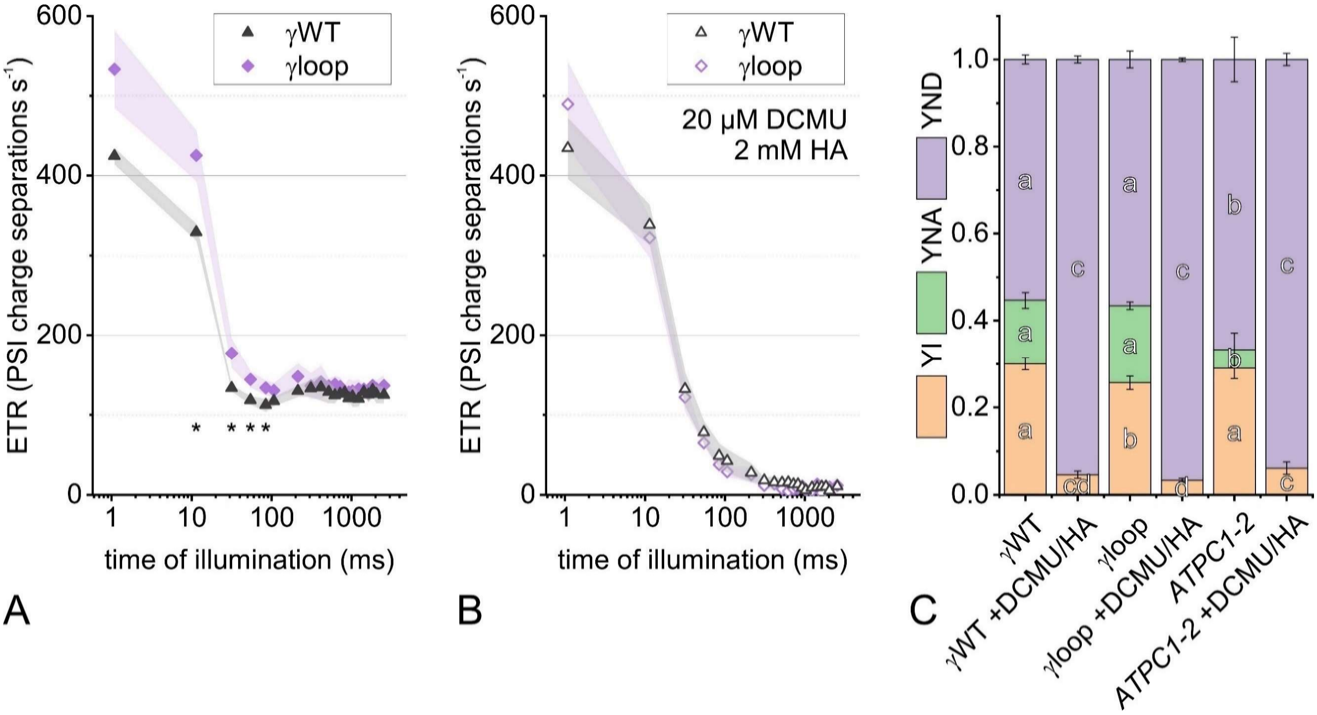


**Fig. S6.** Electron transfer rates (ETR) and PSI redox states during photosynthetic induction treatments are shown. (*A*) Electron transfer developments during a 2.5-s illumination are shown for γWT and γloop ($N$ = 3 ± SD; *Student's t-test, $P$<0.05). PSI-normalized ETR were produced by linear and cyclic electron routes through all photosynthetic complexes. (*B*) shows the samples from panel *A* in the presence of PSII inhibitors, allowing for cyclic electron flow only. (*C*) The redox state quantification at the end of the 2.5-s illumination is shown for $P_{700}$, the primary PSI electron donor. The $P_{700}$ parameters refer to the fractions of photooxidizable (YI; yield of PSI), non-photooxidizable (YNA; acceptor-side limited) and pre-oxidized in actinic light (YND; donor-side limited). The $P_{700}$ analysis also shows the lesion mutant *ATPC1-2* (10) where non-functional $CF_1F_o$ increases YND ($N$ = 3 ± SD, parameter groups analyzed with One-Way ANOVA/Fisher-LSD, $P$<0.05).

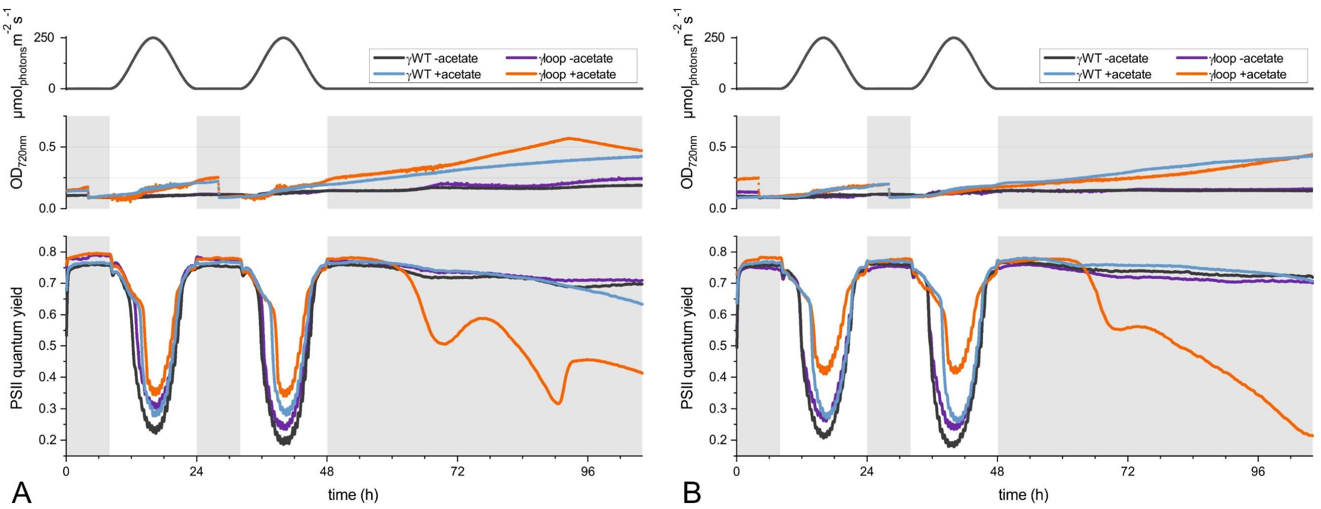


**Fig. S7.** Algal photobioreactor experiments show two light cycles and extended darkness. The $OD_{720nm}$ turbidity, adjusted during the first night cycle (and second in the presence of acetate), as well as the PSII quantum yields are shown for $CF_1F_o$ variants γWT and γloop mutant at a given light intensity. (*A*) and (*B*) show additional transformant lines in support of Main Manuscript Figure 3A.

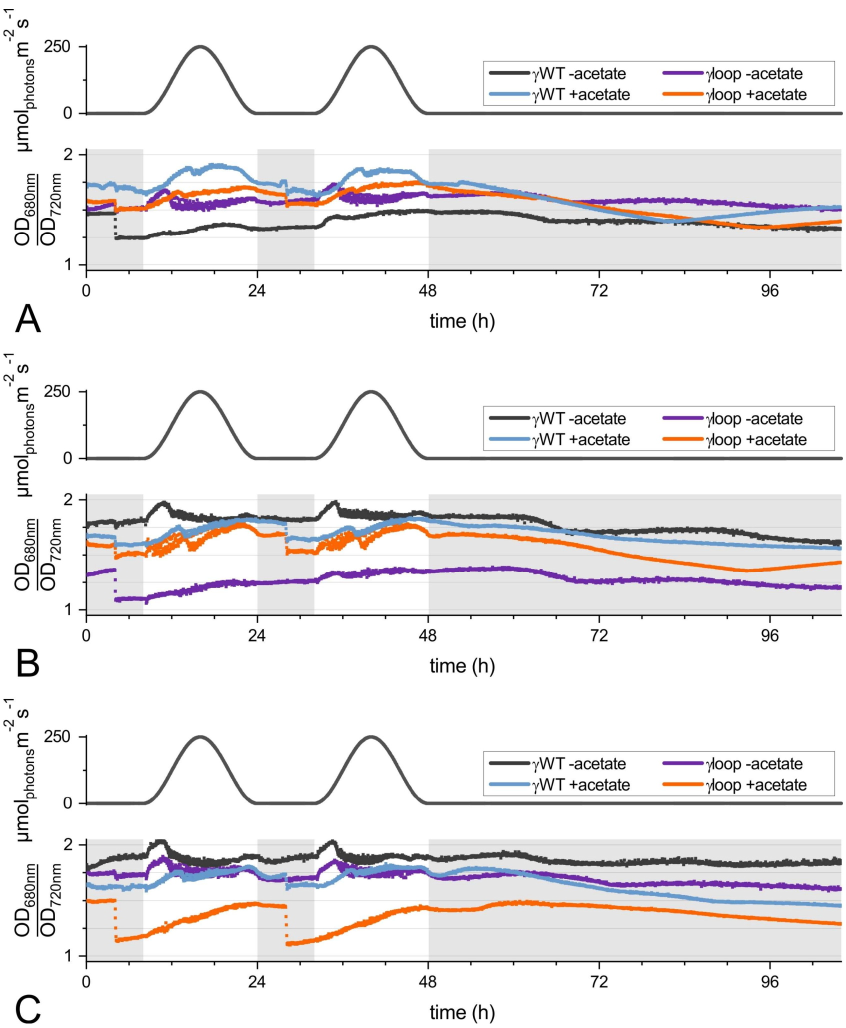


**Fig. S8.** The $OD_{680nm}/OD_{720nm}$ ratios of the algal photobioreactor experiments are shown for $CF_1F_o$ variants γWT and γloop mutant. The ratios are indicative of chlorophyll content and correspond to the experiments shown in (*A*) the Main Manuscript Figure 3A, whereas (*B*) and (*C*) are taken from Figs. S7A and S7B.

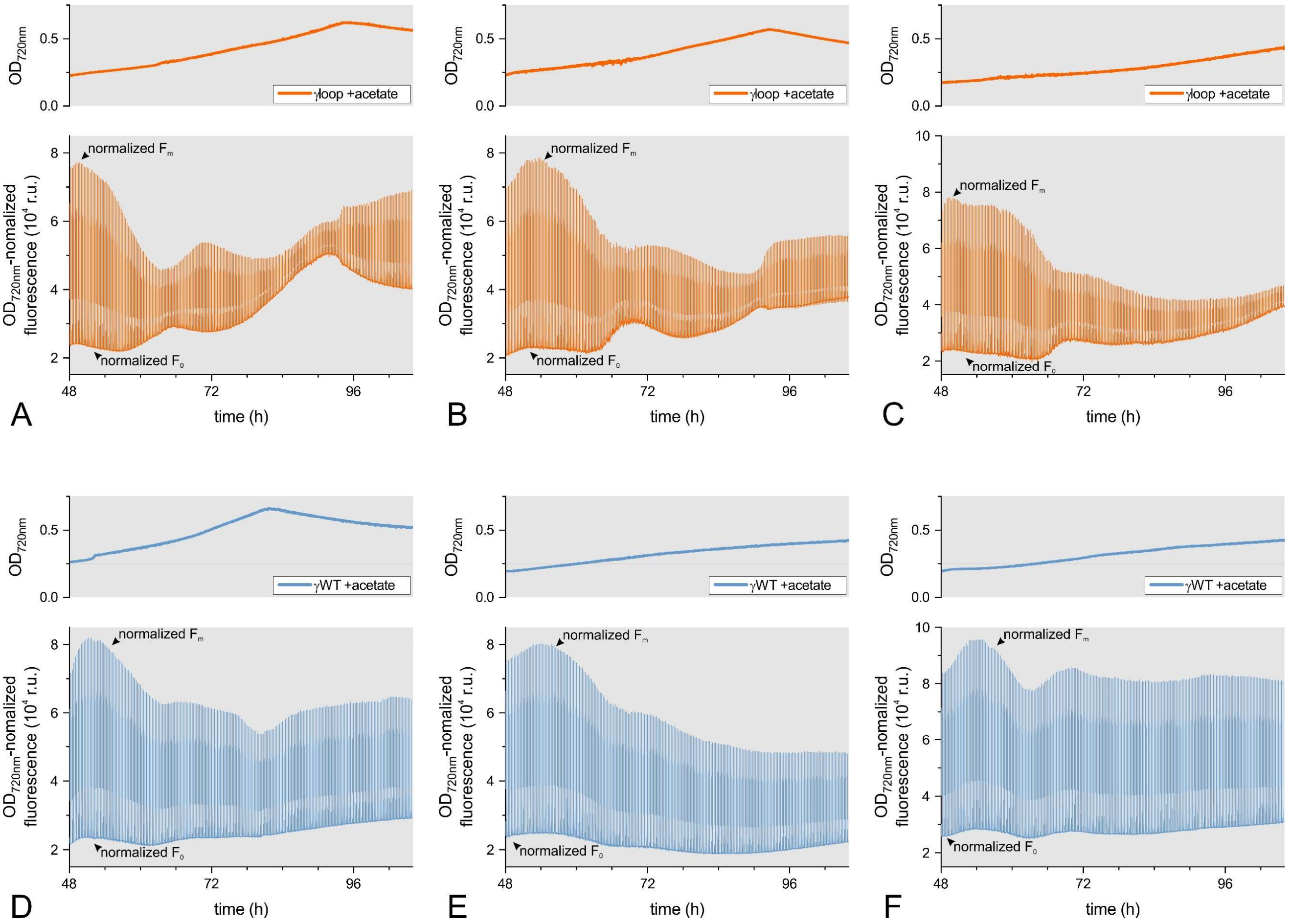


**Fig. S9.** The normalized basal ($F_0$) and maximal chlorophyll fluorescence ($F_m$) is shown in the presence of acetate for three independent transformants of γloop (panels *A*-*C*) and γWT (panels *D*-*F*). The $OD_{720nm}$ values are plotted as well. Panels (*A*) and (*D*) are taken from the Main Manuscript Figure 3A after the second photoperiod. Panels (*B*) and (*E*) are taken from Fig. S7A. Panels (*C*) and (*F*) are taken from Fig. S7B.

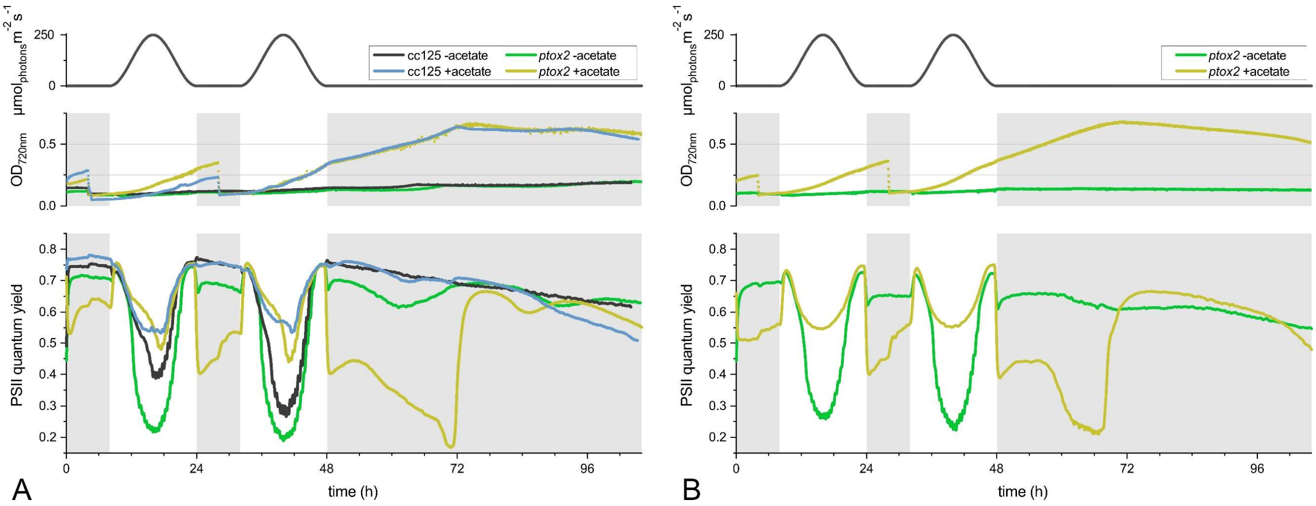


**Fig. S10.** Algal photobioreactor experiments show two light cycles and exposure to extended darkness. (*A*) The $OD_{720nm}$ turbidity, adjusted during the first night cycle (and second in the presence of acetate), as well as the PSII quantum yields are shown for *ptox2* and its reference CC-125. The panel of both plus mating type strains is a biological replicate in support of Main Manuscript Figure 3B. (*B*) The same experiment is shown for an independent *ptox2* strain in the minus mating type background.

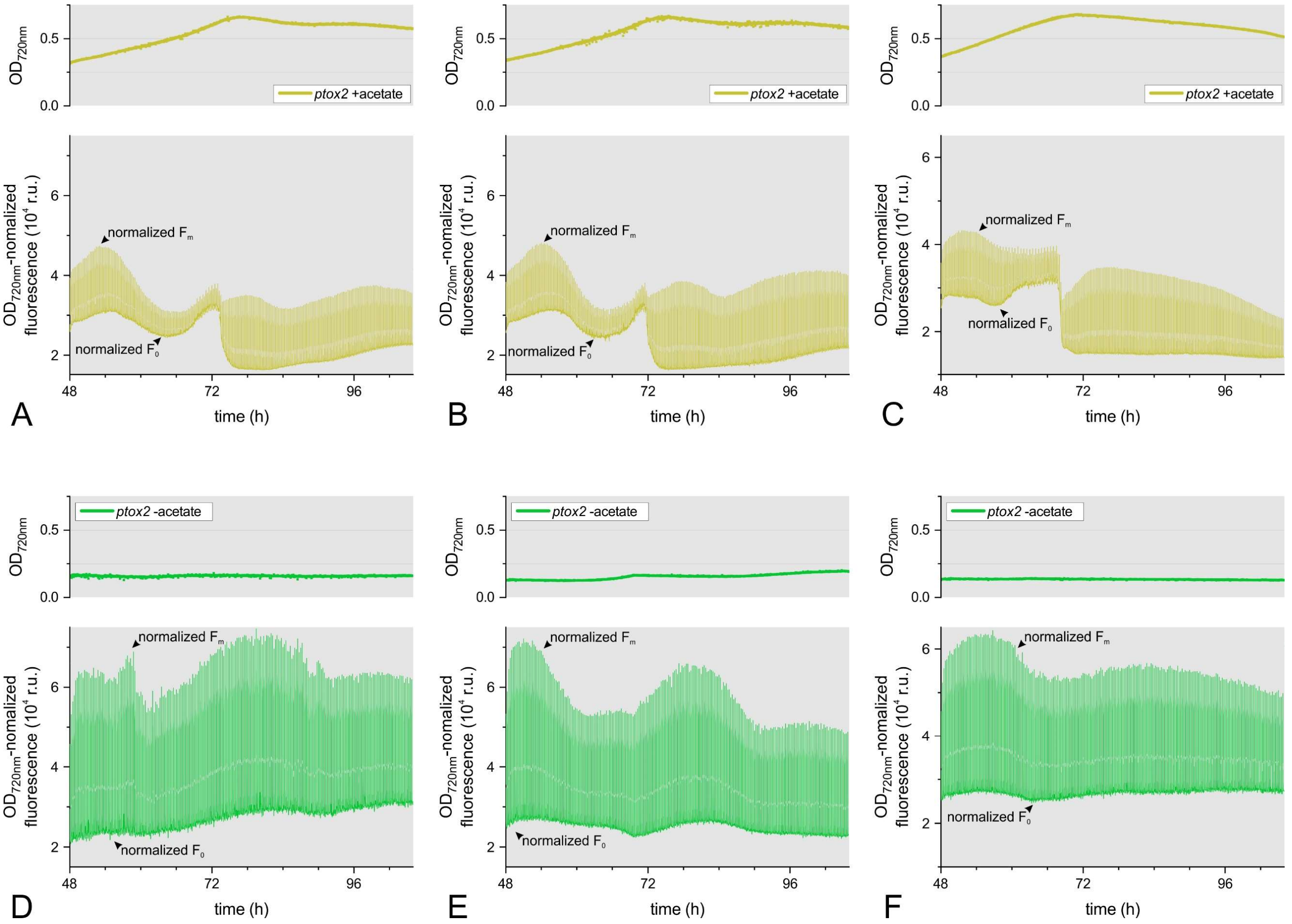


**Fig. S11.** The normalized basal ($F_0$) and maximal chlorophyll fluorescence ($F_m$) is shown in *ptox2* in the presence (panels *A*-*C*) and absence of acetate (panels *D*-*F*). The $OD_{720nm}$ values are plotted as well. Panels (*A*) and (*D*) are taken from the Main Manuscript Figure 3B after the second photoperiod. Panels (*B*) and (*E*) are taken from Fig. S10A. Panels (*C*) and (*F*) are taken from Fig. S10B.

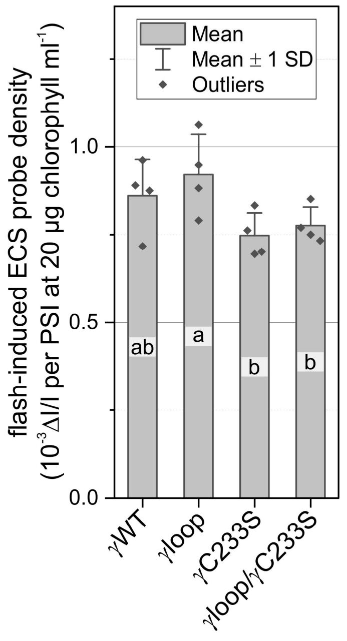


**Fig. S12.** The relationship of optical ECS changes is shown that was produced per PSI turnover and chlorophyll content in the cuvette. The $CF_1F_o$ variants from the Main Manuscript received a saturating 6-ns laser flash in the presence of 1 mM hydroxylamine and 10 µM 3-(3,4-dichlorophenyl)-1,1-dimethylurea to inhibit PSII, and mean signal amplitudes are plotted (520nm – 546nm; $N = 4 \pm$ SD, One-Way ANOVA/Fisher-LSD, $P<0.05$). The plotted ΔI/I points were then used to convert the ECS data in this study into units expressed as PSI charge separations.
